## Supplementary Information for "III. Geometrical framework for thinking about globular proteins: turns in proteins"

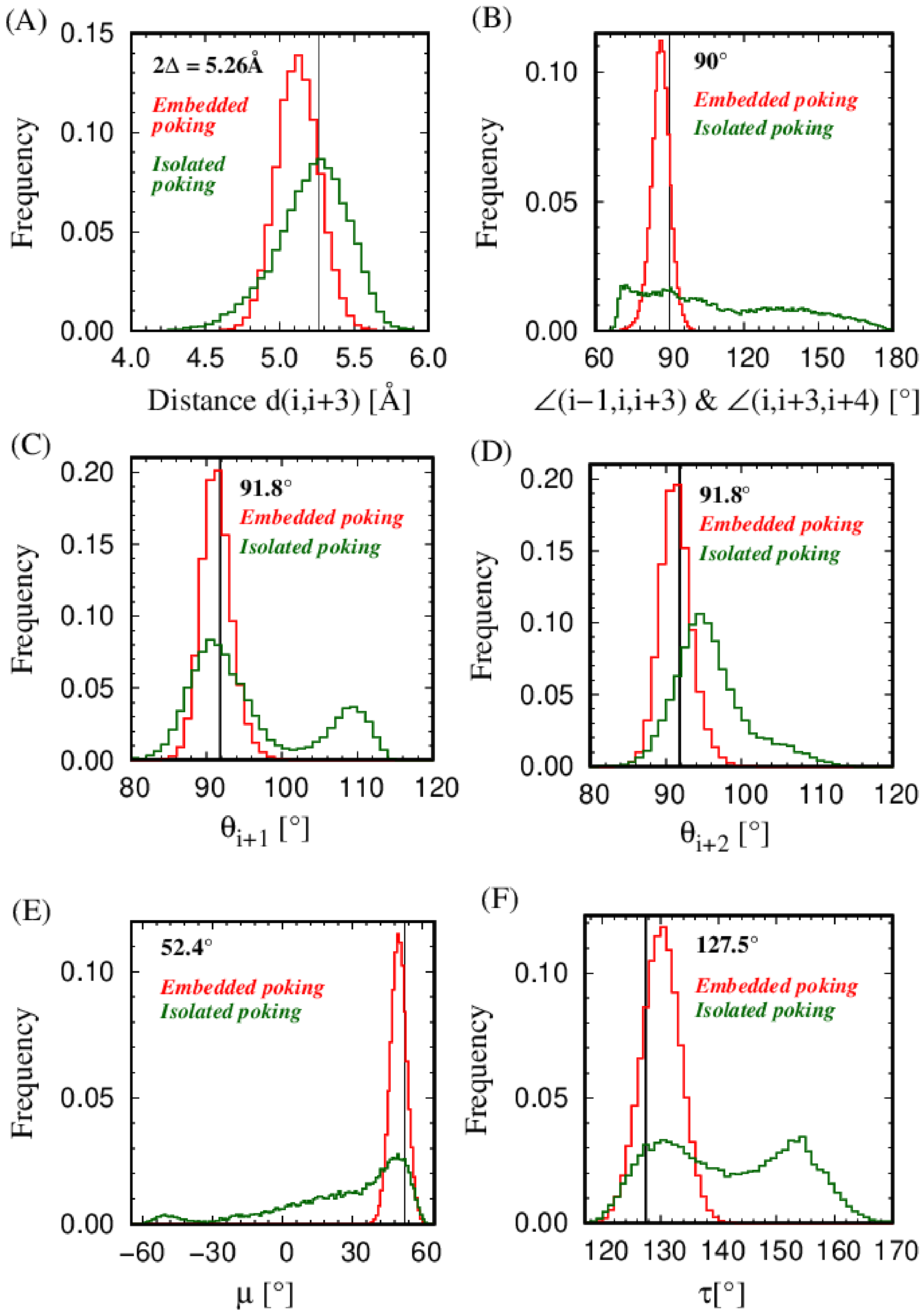

**Figure 1 SI: Geometrical characteristics of embedded and isolated (i,i+3) local poking contacts in our data set comprising 4,391 globular protein chains. There are 176,711 embedded poking contacts for which both the neighboring pairs (i-1,i+2) and (i+1,i+4) are also poking contacts and 21, 571 isolated poking contacts located within protein loops for which neither neighboring pair, (i-1,i+2) or (i+1,i+4), is poking even in an asymmetric manner. An embedded (i,i+3) poking contact is most likely to be within a helix. The red curves depict the histograms for embedded poking contacts, whereas the green curves show the corresponding histograms for isolated poking contacts located within protein loops. The black vertical lines indicate the theoretically predicted values of a given geometrical attribute for the Kepler helix. Panel (A) shows the distribution of the (i,i+3) distances. The black vertical line indicates the theoretically predicted coin diameter of 2Δ = 5.26Å, pertaining to Kepler touching. Panel (B) shows the distribution of the angles (i-1,i,i+3) and (i,i+3,i+4) that are predicted to be 90° (shown as the black vertical line) for embedded poking contacts. Note that although isolated poking contacts in protein loops roughly follow the 2Δ distance constraint of the Kepler helix, they do not necessarily satisfy the 90° angle requirement. Panels (C) and (D) show the distribution of the bond bending angle $\theta_{i+1}$, subtended at point (i+1) by points i and (i+2) and the bond bending angle $\theta_{i+2}$, subtended at point (i+2) by points (i+1) and (i+3 respectively. The black vertical lines in both panels show the ideal value of the bond bend-**

**ing angle of 91.8° in the Kepler helix. Panel (E) shows the distribution of the dihedral angle μ for the quartet of beads (i,i+1,i+2,i+3). This is the angle between the planes formed by [i,(i+1),(i+2)] and [(i+1),(i+2),(i+3)]. The vertical black line indicates the value of 52.4° for the Kepler helix. Panel (F) shows the distribution of the turn angle τ for the quartet of beads (i,i+1,i+2,i+3), defined as the angle between the unit vectors along the directions (i,i+1) and (i+2,i+3). The black vertical line shows the value of 127.5°, the turn angle in the Kepler helix.**

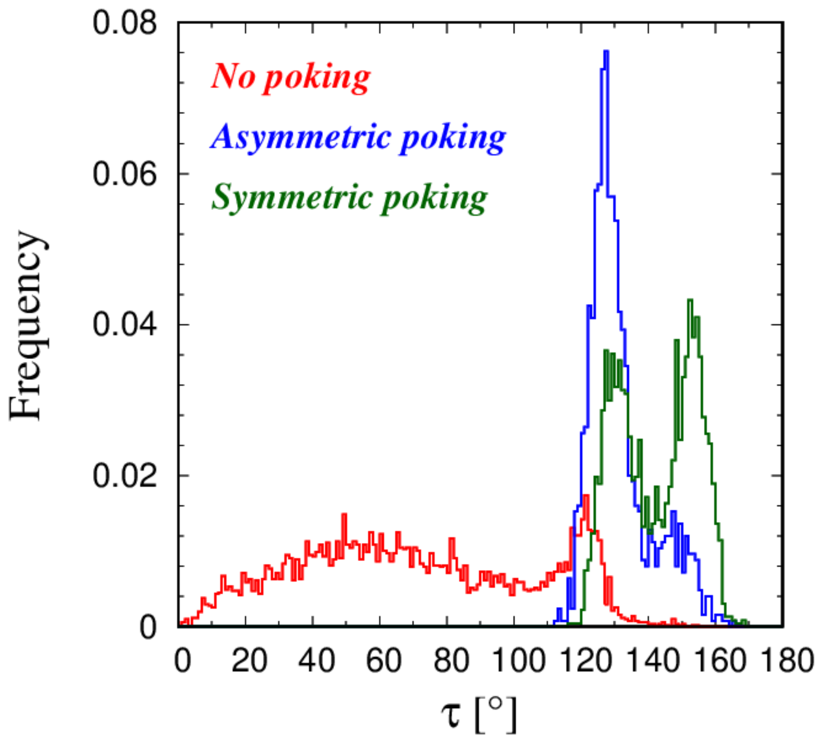


**Figure 2 SI: Frequency distributions of the turn angles τ for three different classes of protein quartets (i,i+1,i+2,i+3). The turn angle τ for a quartet is defined as the angle between the unit vectors along the directions**

**(i,i+1) and (i+2,i+3). Here we have considered the quartets coming from the shortest protein loops of length four for which the quartet itself represents the whole loop. There are 8,563 loops of length four in our data set of 4,391 globular protein chains. The histogram in red represents the frequency distribution of the turn angle $\tau$ for 5,048 quartets that do not display poking interaction of any kind between the beads i and i+3. The histogram in blue shows the frequency distribution of turn angles $\tau$ for 1,247 quartets that display asymmetric or 'one-way' poking contact between beads i and i+3. Finally, the histogram in green is the frequency distribution of the turn angle $\tau$ for 2,268 quartets that have (i,i+3) symmetric poking contacts. Note that quartets with symmetric poking contacts between beads i and i+3 are most effective in changing the chain direction.**

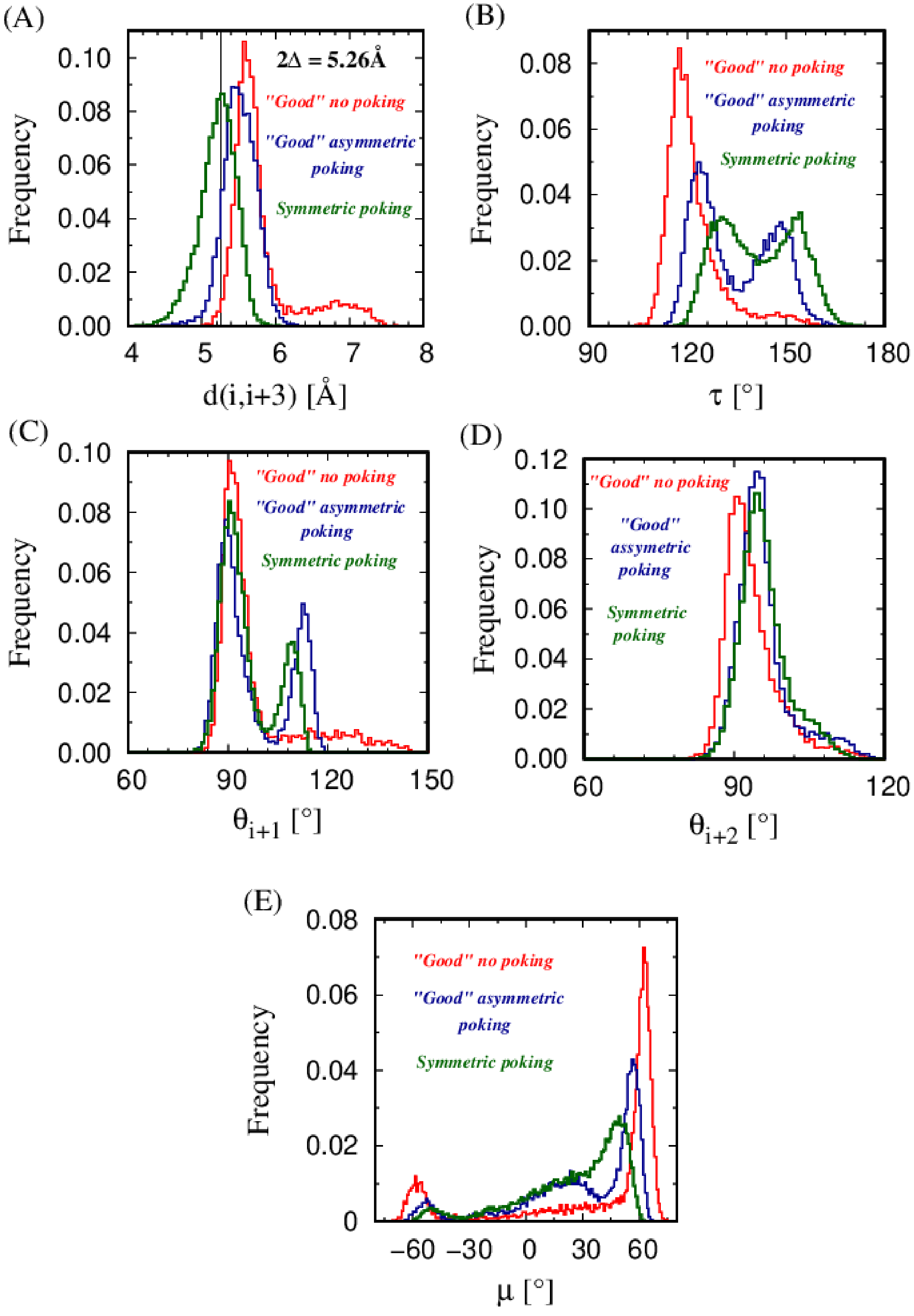
(A)
2Δ = 5.26Å
"Good" no poking
"Good" asymmetric poking
Symmetric poking
Frequency
d(i,i+3) [Å]
(B)
"Good" no poking
"Good" asymmetric poking
Symmetric poking
Frequency
τ [°]
(C)
"Good" no poking
"Good" asymmetric poking
Symmetric poking
Frequency
θi+1 [°]
(D)
"Good" no poking
"Good" assymetric poking
Symmetric poking
Frequency
θi+2 [°]
(E)
"Good" no poking
"Good" asymmetric poking
Symmetric poking
Frequency
μ [°]

**Figure 3 SI: Geometrical characteristics of three types of 'good' isolated local (i,i+3) contacts found in the 59,464 loops of 4,391 globular protein chains. 21,571 isolated symmetric ('two-way') poking (i,i+3) contacts for which neither neighboring pair, (i-1,i+2) or (i+1,i+4), is poking even in an asymmetric manner (green color); 12,297 asymmetric ('one-way') isolated (i,i+3) poking contacts whose 'effective' distance from the symmetric condition is smaller than 0.263Å (5% of 2Δ) (blue color); and finally 6,500 (i,i+3) pairs without any poking contacts between any pairs of beads (i-1,i+2), (i,i+3), or (i+1,i+4) even within a grace distance 0.263Å (5% of 2Δ) (red color). Panel (A) shows the distribution of the (i,i+3) distances. The black vertical line indicates the theoretically predicted coin diameter of 2Δ = 5.26Å, pertaining to Kepler touching. Panel (B) shows the distribution of the turn angle τ. Panel (C) shows the distribution of the bond bending angle $\theta_{i+1}$, the angle subtended at point (i+1) by points i and (i+2). Panel (D) shows the distribution of the bond bending angle $\theta_{i+2}$, the angle subtended at point (i+2) by points (i+1) and (i+3). Panel (E) shows the distribution of the dihedral angle μ, the angle between the planes formed by [i,(i+1),(i+2)] and [(i+1),(i+2),(i+3)], respectively. The geometrical characteristics of the class of 'good' but not fully developed symmetric poking (i,i+3) contacts (those without any poking and those with asymmetric or 'one-way' poking) gradually tend to the geometrical characteristics of the local (i,i+3) contacts in which symmetric poking is fully established (red and blue histograms tend to the green histograms).**

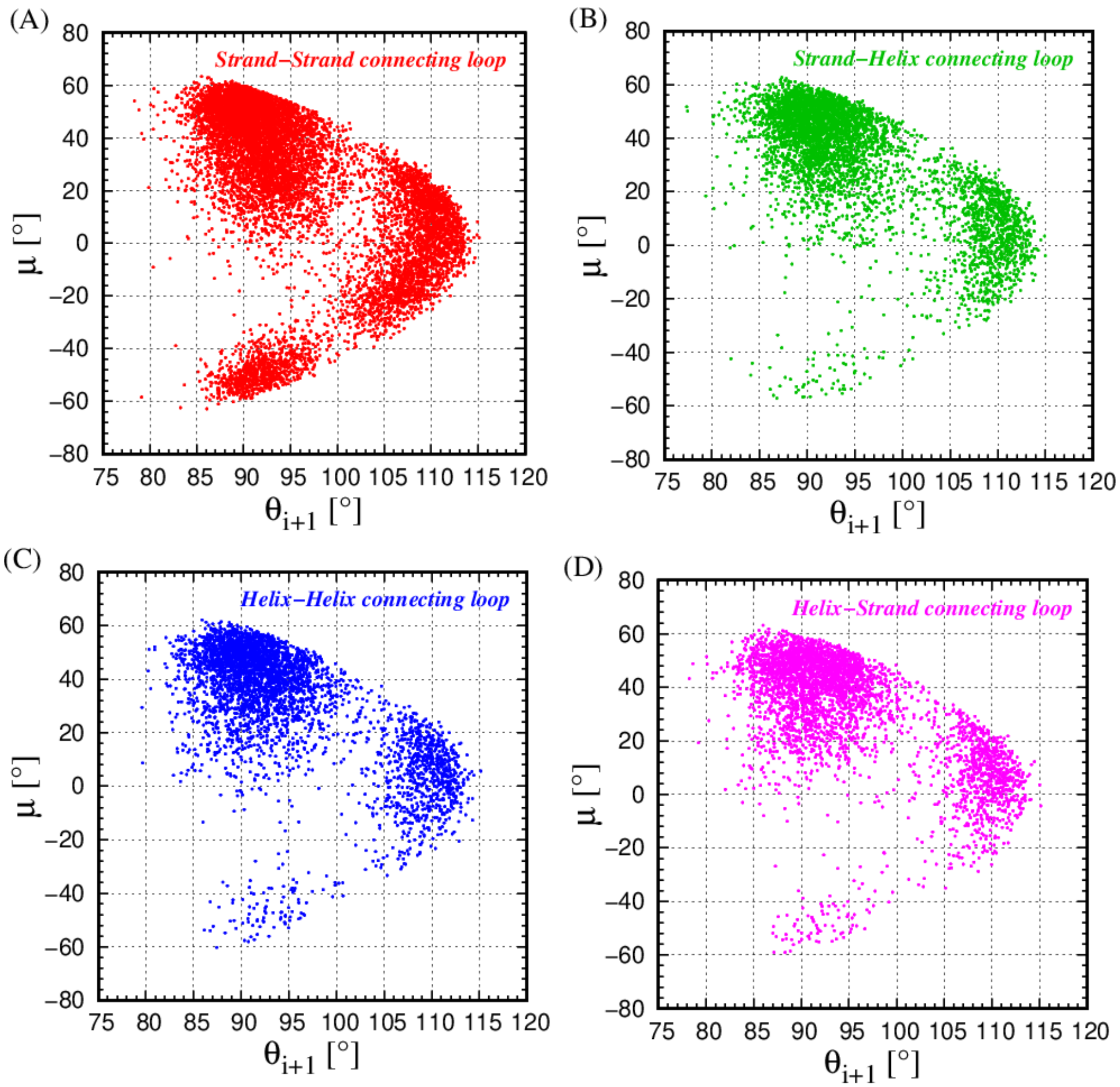


**Figure 4 SI: ($\theta_{i+1}$,μ) cross plots for isolated (i,i+3) symmetric poking contacts located in the loops that connect: a β-strand with a β-strand (9,514 red points) (Panel A); a β-strand with an α-helix (4,493 green points) (Panel B); an α-helix with an α-helix (3,851 blue points) (Panel C); and an α-helix with a β-strand (3,713 magenta points) (Panel D). The cross plots are qualitatively similar underscoring the ability of the same turn types to serve in distinct contexts.**

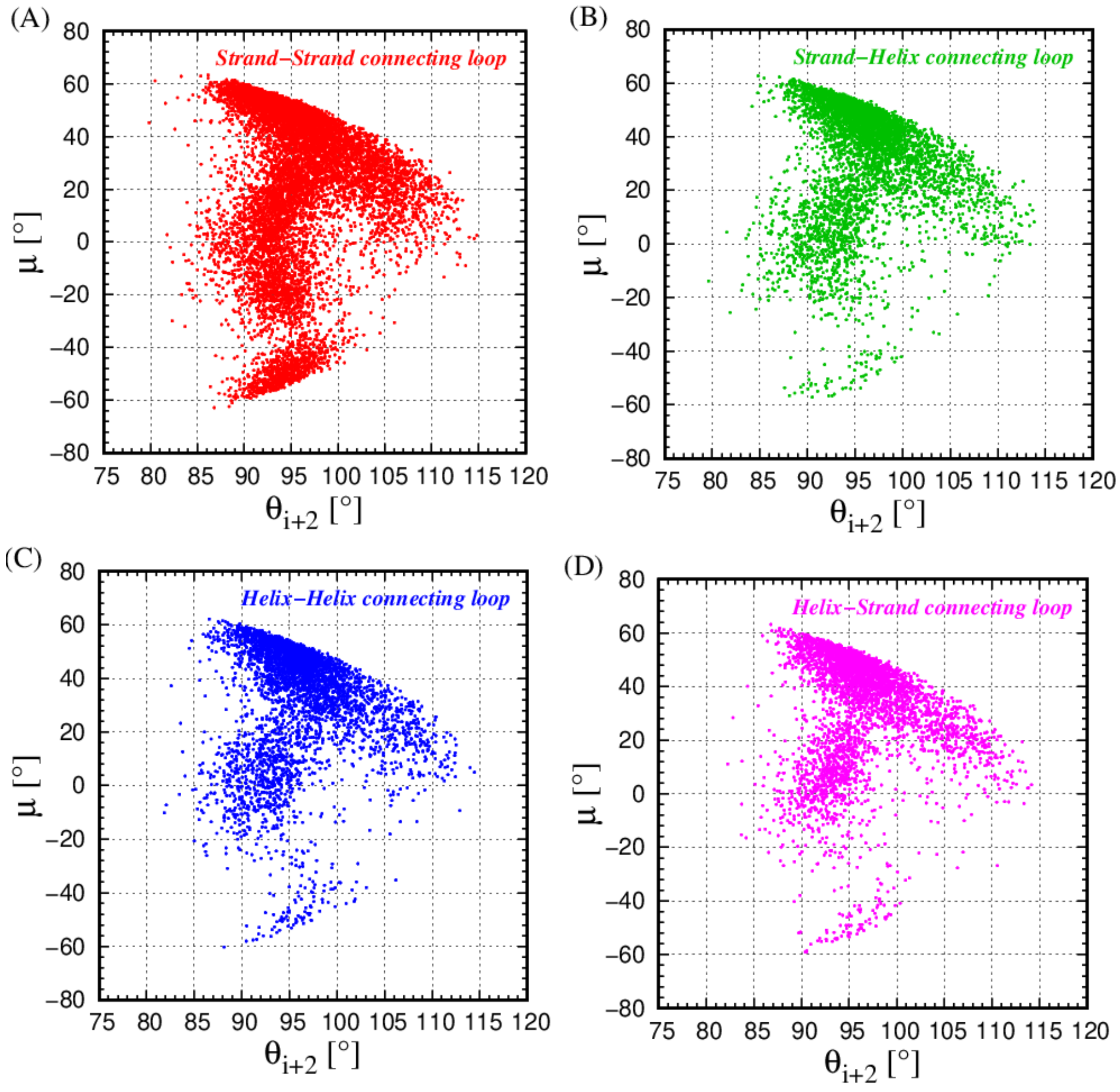


**Figure 5 SI: ($\theta_{i+2}$,μ) cross plots for isolated (i,i+3) symmetric poking contacts located in the loops that connect: a β-strand with a β-strand (9,514 red points) (Panel A); a β-strand with an α-helix (4,493 green points) (Panel B); an α-helix with an α-helix (3,851 blue points) (Panel C); and an α-helix with a β-strand (3,713 magenta points) (Panel D). The cross plots are qualitatively similar underscoring the ability of the same turn types to serve in distinct contexts.**

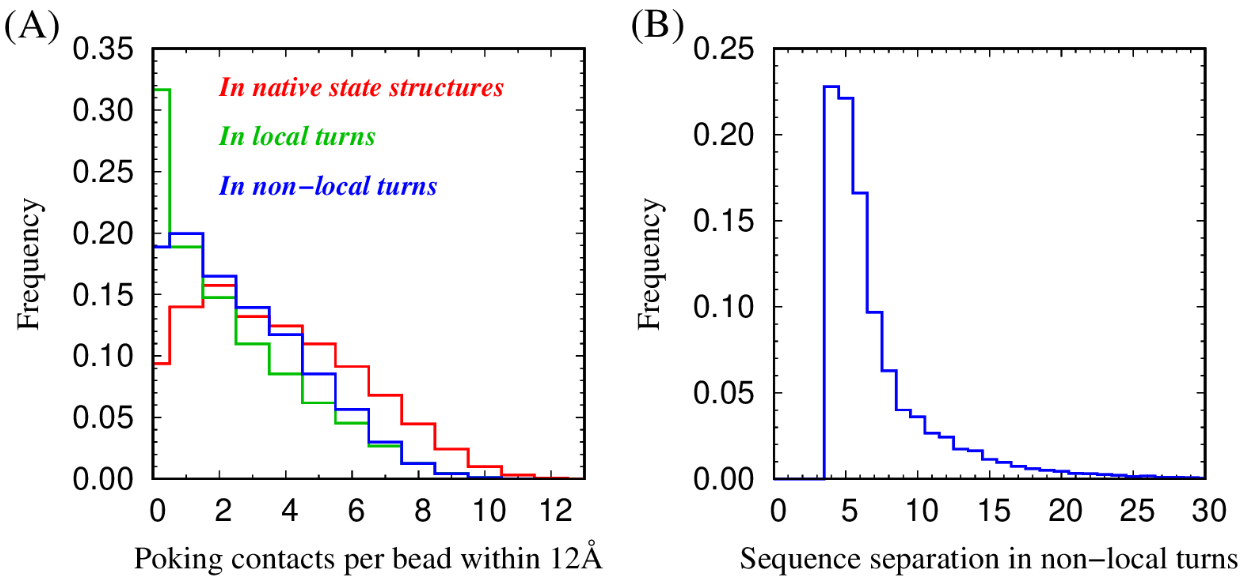


**Figure 6 SI: Panel (A) shows the frequency distribution of the total number of symmetric poking contacts (for all sequence separations along the chain) per bead within 12Å in three different cases: for all beads in the native states of 4,391 proteins of our data set (red histogram); for the beads (i+1) and (i+2) of 21, 571 local turns with (i,i+3) isolated symmetric poking contacts (green histogram); and for the beads (i+1) and (i+2) of 23,115 non-local turns identified by means of (i,j) isolated symmetric poking contacts with j>i+3 (blue histogram). We conclude that beads (i+1) and (i+2) are the most 'orphaned' in (i,i+3) isolated protein turns, meaning that they have the smallest number of poking contacts with other parts of the chain, when compared with the non-local (i,j) non-local turns, j>i+3. Both local and non-local turns have significantly less poking contacts with other parts of the chain than a typical amino acid anywhere in the protein chain (red histogram lies below the green and blue**

**histograms). Panel (B) shows the frequency distribution of the (j-i) sequence separation of the non-local isolated symmetric poking contact (i,j) within a distance of 6Å. Sequence separations of 4 and 5 comprise approximately 50% of all non-local isolated symmetric poking contacts within a distance of 6Å.**

| **Aminoacid type** | **Aminoacid number** | **Aminoacid frequency** |
|---|---|---|
| LEU | 85,661 | 0.0882 |
| ALA | 82,830 | 0.0853 |
| GLY | 76,106 | 0.0784 |
| VAL | 70,032 | 0.0721 |
| GLU | 63,244 | 0.0651 |
| ASP | 57,797 | 0.0595 |
| SER | 56,931 | 0.0586 |
| LYS | 55,756 | 0.0574 |
| ILE | 54,496 | 0.0561 |
| THR | 53,994 | 0.0556 |
| ARG | 47,203 | 0.0486 |
| PRO | 44,768 | 0.0461 |
| ASN | 42,563 | 0.0438 |
| PHE | 39,166 | 0.0403 |
| GLN | 35,078 | 0.0361 |
| TYR | 35,022 | 0.0361 |
| HIS | 22,608 | 0.0233 |
| MET | 19,714 | 0.0203 |
| TRP | 14,704 | 0.0151 |
| CYS | 13,222 | .0136 |

**Table I SI: Total number and frequencies of occurrences of 20 amino acid types in 970,896 residues in our data set of 4,391 globular protein chains.**

### Embedded (i,i+3) poking contacts

| Aminoacids in (i+1) position | Normalized frequency | Aminoacids in (i+2) position | Normalized frequency | Aminoacid pair (i+1,i+2) | Normalized frequency |
|---|---|---|---|---|---|
| ALA | 1.56 | ALA | 1.62 | ALA - ALA | 2.65 |
| GLU | 1.44 | LEU | 1.46 | GLU - ALA | 2.43 |
| LEU | 1.37 | MET | 1.41 | ALA - LEU | 2.40 |
| GLN | 1.34 | GLN | 1.36 | GLN - ALA | 2.28 |
| MET | 1.31 | ARG | 1.32 | ALA - MET | 2.26 |
| ARG | 1.23 | GLU | 1.31 | GLN - GLN | 2.21 |
| ILE | 1.18 | ILE | 1.25 | LEU - ALA | 2.17 |
| LYS | 1.12 | LYS | 1.19 | MET - ALA | 2.15 |
| VAL | 0.99 | PHE | 0.99 | GLU - LEU | 2.12 |
| PHE | 0.95 | VAL | 0.98 | ALA - ARG | 2.08 |
| TRP | 0.92 | TRP | 0.93 | MET - MET | 2.07 |
| TYR | 0.86 | CYS | 0.91 | LYS - ALA | 2.05 |
| CYS | 0.86 | TYR | 0.88 | ARG - ALA | 2.04 |
| ASP | 0.84 | HIS | 0.79 | GLU - GLU | 2.04 |
| HIS | 0.81 | ASP | 0.74 | LEU - ARG | 2.04 |
| THR | 0.74 | THR | 0.69 | GLN - LEU | 2.03 |
| ASN | 0.68 | ASN | 0.68 | GLU - MET | 2.01 |
| SER | 0.67 | SER | 0.64 | ALA- ILE | 2.00 |
| GLY | 0.43 | GLY | 0.36 | LEU - GLU | 1.99 |
| PRO | 0.17 | PRO | 0.02 | ALA - GLN | 1.98 |

**Table II SI: Normalized frequencies of occurrence of 20 amino acid types in the positions (i+1) and (i+2) of 176,711 (i,i+3) embedded symmetric poking contacts, as well as the top 20 over-expressed (i+1,i+2) amino acid pairs.**

### Isolated (i,i+3) poking contacts: Region I

| Aminoacids in (i+1) position | Normalized frequency | Aminoacids in (i+2) position | Normalized frequency | Aminoacid pair (i+1,i+2) | Normalized frequency |
|---|---|---|---|---|---|
| PRO | 3.37 | ASP | 2.86 | PRO - ASP | 10.86 |
| SER | 1.63 | ASN | 2.45 | PRO - ASN | 8.83 |
| GLU | 1.48 | HIS | 1.51 | LYS - ASP | 6.23 |
| LYS | 1.37 | TRP | 1.38 | PRO - HIS | 4.94 |
| ALA | 1.30 | TYR | 1.32 | PRO - GLU | 4.87 |
| ASP | 1.15 | SER | 1.24 | PRO - TRP | 4.80 |
| ARG | 1.00 | GLN | 1.19 | ALA - ASP | 4.38 |
| GLN | 0.97 | LYS | 1.16 | PRO - SER | 4.30 |
| TRP | 0.89 | PHE | 1.13 | GLU - ASP | 4.24 |
| CYS | 0.80 | GLU | 1.12 | PRO - THR | 4.08 |
| HIS | 0.71 | THR | 0.92 | LYS - ASN | 4.02 |
| THR | 0.70 | ARG | 0.84 | PRO - GLN | 3.95 |
| ASN | 0.68 | CYS | 0.77 | SER - ASP | 3.75 |
| TYR | 0.62 | LEU | 0.68 | GLU - ASN | 3.70 |
| LEU | 0.60 | MET | 0.66 | GLN- HIS | 3.46 |
| MET | 0.59 | ALA | 0.62 | PRO - TYR | 3.40 |
| PHE | 0.58 | GLY | 0.52 | ASP - ASP | 3.37 |
| ILE | 0.48 | ILE | 0.30 | ALA - ASN | 3.32 |
| GLY | 0.45 | VAL | 0.29 | SER - ASN | 3.31 |
| VAL | 0.42 | PRO | 0.22 | TRP - TRP | 3.21 |

**Table III SI: Normalized frequencies of occurrence of 20 amino acid types in the positions (i+1) and (i+2) of 9,604 local (i,i+3) isolated symmetric poking contacts belonging to Region I, as well as the top 20 over-expressed (i+1,i+2) amino acid pairs.**

### Isolated (i,i+3) poking contacts: Region II

| Aminoacids in (i+1) position | Normalized frequency | Aminoacids in (i+2) position | Normalized frequency | Aminoacid pair (i+1,i+2) | Normalized frequency |
|---|---|---|---|---|---|
| PRO | 1.89 | ASN | 2.46 | PRO - ASN | 5.44 |
| GLU | 1.69 | ASP | 1.99 | PRO - ASP | 5.33 |
| LYS | 1.62 | THR | 1.74 | LYS - TYR | 4.95 |
| ASP | 1.41 | HIS | 1.68 | LYS - ASP | 4.55 |
| SER | 1.40 | TYR | 1.62 | GLU - ASN | 4.52 |
| ALA | 1.29 | PHE | 1.48 | LYS - ASN | 4.36 |
| GLN | 1.16 | TRP | 1.13 | GLU - HIS | 3.99 |
| ARG | 1.02 | SER | 1.11 | SER - CYS | 3.55 |
| THR | 0.92 | CYS | 1.05 | PRO - THR | 3.48 |
| HIS | 0.86 | ARG | 1.03 | ALA - ASN | 3.45 |
| ASN | 0.80 | LYS | 0.94 | SER - ASN | 3.38 |
| TRP | 0.73 | GLU | 0.81 | ARG - TYR | 3.37 |
| TYR | 0.68 | GLN | 0.80 | GLN - TYR | 3.36 |
| VAL | 0.61 | ILE | 0.78 | GLU - THR | 3.36 |
| LEU | 0.60 | VAL | 0.69 | PRO - HIS | 3.35 |
| MET | 0.60 | GLY | 0.59 | TRP - HIS | 3.23 |
| CYS | 0.59 | MET | 0.56 | ARG - HIS | 3.01 |
| ILE | 0.48 | LEU | 0.54 | HIS - ASP | 3.00 |
| GLY | 0.47 | ALA | 0.40 | LYS - THR | 2.97 |
| PHE | 0.45 | PRO | 0.09 | GLU - TYR | 2.92 |

**Table IV SI: Normalized frequencies of occurrence of 20 amino acid types in the positions (i+1) and (i+2) of 5,271 local (i,i+3) isolated symmetric poking contacts belonging to Region II, as well as the top 20 over-expressed (i+1,i+2) amino acid pairs.**

### Isolated (i,i+3) poking contacts: Region III

| Aminoacids in (i+1) position | Normalized frequency | Aminoacids in (i+2) position | Normalized frequency | Aminoacid pair (i+1,i+2) | Normalized frequency |
|---|---|---|---|---|---|
| PRO | 3.67 | GLY | 9.02 | PRO - GLY | 34.25 |
| LYS | 1.79 | ASN | 1.50 | GLU - GLY | 17.06 |
| GLU | 1.63 | ASP | 0.79 | LYS - GLY | 16.85 |
| ASP | 1.04 | CYS | 0.57 | ALA - GLY | 9.87 |
| ARG | 1.03 | HIS | 0.52 | ARG - GLY | 9.81 |
| ALA | 1.02 | TYR | 0.41 | ASP - GLY | 9.09 |
| GLN | 0.95 | PHE | 0.38 | GLN - GLY | 8.94 |
| SER | 0.91 | ARG | 0.37 | VAL - GLY | 8.88 |
| HIS | 0.86 | GLN | 0.34 | SER - GLY | 8.28 |
| VAL | 0.85 | SER | 0.28 | HIS - GLY | 7.59 |
| ASN | 0.70 | LYS | 0.27 | ILE - GLY | 7.26 |
| ILE | 0.69 | GLU | 0.23 | PRO - ASN | 5.81 |
| TYR | 0.65 | THR | 0.21 | ASN - GLY | 5.76 |
| THR | 0.58 | ALA | 0.20 | MET - CYS | 5.42 |
| MET | 0.57 | TRP | 0.18 | THR - GLY | 5.18 |
| TRP | 0.55 | MET | 0.16 | TYR - GLY | 5.15 |
| GLY | 0.54 | LEU | 0.11 | PHE - GLY | 3.92 |
| PHE | 0.49 | VAL | 0.05 | LEU - GLY | 3.71 |
| LEU | 0.47 | PRO | 0.04 | LYS - ASN | 3.68 |
| CYS | 0.18 | ILE | 0.03 | TRP - GLY | 3.40 |

**Table V SI: Normalized frequencies of occurrence of 20 amino acid types in the positions (i+1) and (i+2) of 3,237 local (i,i+3) isolated symmetric poking contacts belonging to Region III, as well as the top 20 over-expressed (i+1,i+2) amino acid pairs.**

### Isolated (i,i+3) poking contacts: Region IV

| Aminoacids in (i+1) position | Normalized frequency | Aminoacids in (i+2) position | Normalized frequency | Aminoacid pair (i+1,i+2) | Normalized frequency |
|---|---|---|---|---|---|
| GLY | 4.31 | ASN | 3.46 | GLY - ASP | 15.79 |
| PRO | 1.64 | GLY | 2.85 | GLY - ASN | 12.03 |
| LYS | 1.13 | ASP | 2.74 | PRO - GLY | 9.02 |
| ASP | 1.00 | CYS | 1.28 | GLY - SER | 8.66 |
| HIS | 0.95 | SER | 1.26 | GLY - THR | 6.30 |
| TYR | 0.91 | THR | 0.82 | LYS - ASP | 5.60 |
| ASN | 0.90 | GLU | 0.80 | HIS - CYS | 5.48 |
| GLU | 0.86 | LYS | 0.77 | LYS - ASN | 5.18 |
| TRP | 0.84 | HIS | 0.73 | GLY - GLU | 4.94 |
| GLN | 0.78 | TRP | 0.72 | LYS - GLY | 4.73 |
| PHE | 0.71 | ARG | 0.68 | GLY - LYS | 4.72 |
| ARG | 0.68 | TYR | 0.54 | TYR - CYS | 4.45 |
| MET | 0.62 | GLN | 0.53 | ASN - ASN | 4.31 |
| ALA | 0.57 | ALA | 0.50 | PRO - ASN | 4.30 |
| SER | 0.49 | PHE | 0.45 | GLY - PRO | 4.21 |
| THR | 0.48 | MET | 0.42 | GLU - ASN | 4.12 |
| LEU | 0.46 | PRO | 0.33 | GLY - ARG | 4.11 |
| CYS | 0.42 | LEU | 0.30 | ASP - GLY | 4.10 |
| VAL | 0.40 | VAL | 0.15 | ASP - ASN | 4.01 |
| ILE | 0.29 | ILE | 0.12 | TYR - ASN | 3.87 |

**Table VI SI: Normalized frequencies of occurrence of 20 amino acid types in the positions (i+1) and (i+2) of 2,298 local (i,i+3) isolated symmetric poking contacts belonging to Region IV, as well as the top 20 over-expressed (i+1,i+2) amino acid pairs.**

### Isolated (i,i+3) poking contacts: Region V

| Aminoacids in (i+1) position | Normalized frequency | Aminoacids in (i+2) position | Normalized frequency | Aminoacid pair (i+1,i+2) | Normalized frequency |
|---|---|---|---|---|---|
| ASN | 5.45 | GLY | 10.24 | ASN - GLY | 57.42 |
| ASP | 3.49 | ASN | 1.32 | ASP - GLY | 34.40 |
| GLY | 1.78 | ASP | 0.57 | GLY - GLY | 17.94 |
| HIS | 1.40 | GLN | 0.29 | LYS - GLY | 13.94 |
| LYS | 1.33 | TRP | 0.28 | HIS - GLY | 13.17 |
| GLN | 1.02 | SER | 0.27 | ARG - GLY | 10.63 |
| GLU | 1.00 | TYR | 0.22 | GLU - GLY | 9.95 |
| ARG | 0.98 | LYS | 0.20 | GLN - GLY | 9.42 |
| ALA | 0.64 | PHE | 0.19 | ASN - ASN | 7.18 |
| SER | 0.54 | CYS | 0.19 | ALA - GLY | 6.56 |
| MET | 0.41 | GLU | 0.18 | SER - GLY | 6.15 |
| TYR | 0.31 | HIS | 0.18 | ASP - ASN | 4.96 |
| PHE | 0.30 | ARG | 0.17 | MET - GLY | 4.20 |
| TRP | 0.23 | THR | 0.12 | HIS - TYR | 4.11 |
| LEU | 0.21 | ALA | 0.06 | ASN - TRP | 3.91 |
| CYS | 0.19 | LEU | 0.05 | TYR - GLY | 3.67 |
| VAL | 0.10 | VAL | 0.04 | GLN - CYS | 3.50 |
| THR | 0.08 | ILE | 0.03 | PHE - GLY | 3.28 |
| PRO | 0.02 | PRO | 0 | ASN - GLN | 3.26 |
| ILE | 0.01 | MET | 0 | GLU - ASN | 2.42 |

**Table VII SI: Normalized frequencies of occurrence of 20 amino acid types in the positions (i+1) and (i+2) of 1,161 local (i,i+3) isolated symmetric poking contacts belonging to Region V, as well as the top 20 over-expressed (i+1,i+2) amino acid pairs.**

### Isolated (i,i+3) poking contacts with 'short' bonds

| **Aminoacids in (i+1) position** | **Normalized frequency** | **Aminoacids in (i+2) position** | **Normalized frequency** | **Aminoacid pair (i+1,i+2)** | **Normalized frequency** |
|---|---|---|---|---|---|
| PRO | 4.089 | PRO | 16.16 | ASN - PRO | 84.63 |
| ASN | 4.00 | ARG | 0.54 | ASP - PRO | 36.81 |
| ASP | 1.77 | TYR | 0.49 | CYS - PRO | 27.95 |
| SER | 1.41 | GLY | 0.45 | HIS - PRO | 24.43 |
| HIS | 1.31 | PHE | 0.44 | MET - PRO | 22.71 |
| CYS | 1.29 | GLU | 0.40 | GLN - PRO | 21.05 |
| MET | 1.26 | HIS | 0.38 | SER - PRO | 20.96 |
| GLN | 1.09 | SER | 0.37 | GLU - PRO | 18.98 |
| VAL | 0.92 | VAL | 0.31 | VAL - PRO | 18.52 |
| GLU | 0.88 | ASN | 0.30 | LYS - PRO | 14.87 |
| ARG | 0.72 | ASP | 0.30 | THR - PRO | 13.70 |
| LYS | 0.69 | TRP | 0.29 | ARG - PRO | 13.70 |
| THR | 0.63 | GLN | 0.24 | TRP - PRO | 12.61 |
| TRP | 0.58 | MET | 0.21 | PRO - PRO | 12.37 |
| ILE | 0.47 | ALA | 0.20 | PRO - ARG | 11.74 |
| ALA | 0.31 | THR | 0.16 | PRO - TYR | 10.58 |
| LEU | 0.20 | LEU | 0.05 | PRO - GLU | 8.76 |
| GLY | 0.17 | LYS | 0 | ILE - PRO | 8.50 |
| TYR | 0.12 | ILE | 0 | PRO - PHE | 7.09 |
| PHE | 0.11 | CYS | 0 | PRO - TRP | 6.30 |

**Table VIII SI: Normalized frequencies of occurrence of 20 amino acid types in the positions (i+1) and (i+2) of 228 protein quartets (i,i+1,i+2,i+3) located in protein loops that have at least one short bond with an isolated symmetric poking contact established between beads i and i+3, along with the top 20 over-expressed (i+1,i+2) amino acid pairs. In 75% of all cases the short bond in question is (i+1,i+2).**
